# O-GlcNAc modification of nuclear pore complexes accelerates bi-directional transport

**DOI:** 10.1101/2020.10.09.334029

**Authors:** Tae Yeon Yoo, Timothy J Mitchison

**Author notes:** Corresponding author:, Address: 200 Longwood Ave., Boston, MA 02115.

## Abstract

Macromolecular transport across the nuclear envelope depends on facilitated diffusion through nuclear pore complexes (NPCs). The interior of NPCs contains a permeability barrier made of phenylalanine-glycine (FG) repeat domains that selectively facilitates the permeation of cargoes bound to nuclear transport receptors (NTRs). FG repeats in NPC are a major site of O-linked N-acetylglucosamine (O-GlcNAc) modification, but the functional role of this modification in nucleocytoplasmic transport is unclear. We developed high-throughput assays based on optogenetic probes to quantify the kinetics of nuclear import and export in living human cells. We found that increasing O-GlcNAc modification of the NPC accelerated NTR-facilitated nucleocytoplasmic transport of proteins in both directions, and decreasing modification slowed transport. Super-resolution imaging revealed strong enrichment of O-GlcNAc at the FG-repeat barrier. O-GlcNAc modification also accelerated passive permeation of a small, inert protein through NPCs. We conclude that O-GlcNAc modification accelerates nucleocytoplasmic transport by enhancing the non-specific permeability the FG-repeat barrier, perhaps by steric inhibition of interactions between FG repeats.

**Summary:** Nuclear pore complexes mediate nuclear transport and are highly modified with O-linked N-acetylglucosamine (O-GlcNAc) on FG repeat domains. Using a new quantitative live-cell imaging assay, Yoo and Mitchison demonstrate acceleration of nuclear import and export by O-GlcNAc modification.

## Introduction

Nucleocytoplasmic transport of macromolecules occurs through thousands of nuclear pore complexes (NPCs) embedded in the nuclear envelope (Callan and Tomlin, 1950; Maul et al., 1972; Watson, 1959). Each NPC consists of ~30 different nucleoporins (NUPs), many of which are FG-NUPs that project intrinsically disordered regions containing phenylalanine-glycine (FG) repeats into the central channel (Denning et al., 2003; Kim et al., 2018; Lin and Hoelz, 2019). The FG domains constitute a selective permeability barrier that allows traffic of molecules in a size-dependent manner (Feldherr and Akin, 1997; Keminer and Peters, 1999). Most large molecules are restricted from permeating through the barrier, unless they are complexed with nuclear transport receptors (NTRs), such as importin-βs and exportins (Fried and Kutay, 2003; Ribbeck and Gorlich, 2001). The transport of large NTR-cargo complex is facilitated by specific, transient interactions between FG repeats and hydrophobic pockets of NTR (Hough et al., 2015; Milles et al., 2015). GTP-bound form of small GTPase Ran (RanGTP) forms concentration gradient across the nuclear envelope and controls the stability of NTR-cargo interactions, thereby dictating the directionality of NTR-facilitated cargo transport (Christie et al., 2016; Macara, 2001; Matsuura, 2016). In this picture, the overall kinetics of nucleocytoplasmic transport is governed by interactions between FG repeats that determine the passive diffusion properties of NPCs and by interactions between NTRs and FG repeats that determine facilitated diffusion.

FG-NUPs are heavily modified by O-linked β-N-acetylglucosamine (O-GlcNAc), where the monosaccharide is reversibly attached to the hydroxyl oxygen of serine and threonine (Davis and Blobel, 1987; Hanover et al., 1987; Holt et al., 1987; Li and Kohler, 2014; Snow et al., 1987). FG-NUPs were some of the first identified O-GlcNAc-modification substrates and are still among the most heavily modified of all known substrates, yet the functional role of this modification is still unclear. A single pair of enzymes, O-GlcNAc transferase (OGT) and O-GlcNAcase (OGA), regulate O-GlcNAcylation of over a thousand of proteins, making it difficult to determine how changes in cell physiology are caused by modification of specific proteins or assemblies (Chatham et al., 2020; Kreppel et al., 1997; Lubas et al., 1997; Wells et al., 2002). Previous *in vitro* studies have shown that O-GlcNAcylation alters the structure and permeability of hydrogels derived from FG domains (Labokha et al., 2013) as well as the radius of gyration of FG domains in solution (Tan et al., 2018). These observations made in simplified biochemical model system predict that O-GlcNAcylation might increase the permeability of NPCs and promote nucleocytoplasmic transport in living cells, but this has never been tested, in large part due to the absence of quantitative tools to measure transport rates in cells. Here, we overcome this limitation by developing optogenetic-based nuclear transport assay and test whether O-GlcNAcylation of the NPC modulates the nuclear transport rates. Alteration of nucleocytoplasmic transport has been implicated in many physiological and pathological processes (Cho and Hetzer, 2020; Grima et al., 2017; Khan et al., 2020; Kim and Taylor, 2017; Lord et al., 2015; Rempel et al., 2019), so tools to measure and modulate it will find multiple uses. In cells, O-GlcNAcylation was proposed to affect NPC integrity and composition (Mizuguchi-Hata et al., 2013; Zhu et al., 2016). This predicts a change in NPC density following O-GlcNAc perturbation, which we also tested.

## Results and Discussion

### Optogenetic-based high-throughput assays are developed to quantify nuclear transport rates in living human cells

To quantify nuclear import and export kinetics we repurposed the NES-mCherry-LINuS and NLS-mCherry-LEXY probes previously created for light-controlled gene expression (Niopek et al., 2014; Niopek et al., 2016). They consist of LINuS (LOV2-based photoactivatable NLS) (Niopek et al., 2014) and LEXY (LOV2-based photoactivatable NES) (Niopek et al., 2016) attached to mCherry along with a constitutive NES or NLS that counteracts the photoactivatable signals (Fig. 1 A, and B). The NES-mCherry-LINuS import probe is mostly localized in the cytoplasm in the dark state but translocates to the nucleus upon blue-light stimulation as the activated LINuS overpowers the NES (Fig. 1A). Likewise, the NLS-mCherry-LEXY export probe translocates mostly from the nucleus to the cytoplasm upon blue-light stimulation (Fig. 1B). The kinetics of light-induced changes in the nuclear intensity of the probes were well-described by mono-exponential decay models. This allowed accurate fitting of kinetic data to output single decay constants which report transport rates. The light-induced conformational change of LOV2 is so fast, within a millisecond (Konold et al., 2016), that it does not contribute to the overall translocation kinetics. The dark-reversibility of LINuS and LEXY allows repeated measurements of the transport rates in individual cells. We generated U2OS cell lines stably expressing the transport probes and automated image acquisitions and analysis to allow high-throughput, time-course measurement and accurate detection of rate changes with high statistical confidence (Fig. 1C).

**Fig. 1.**
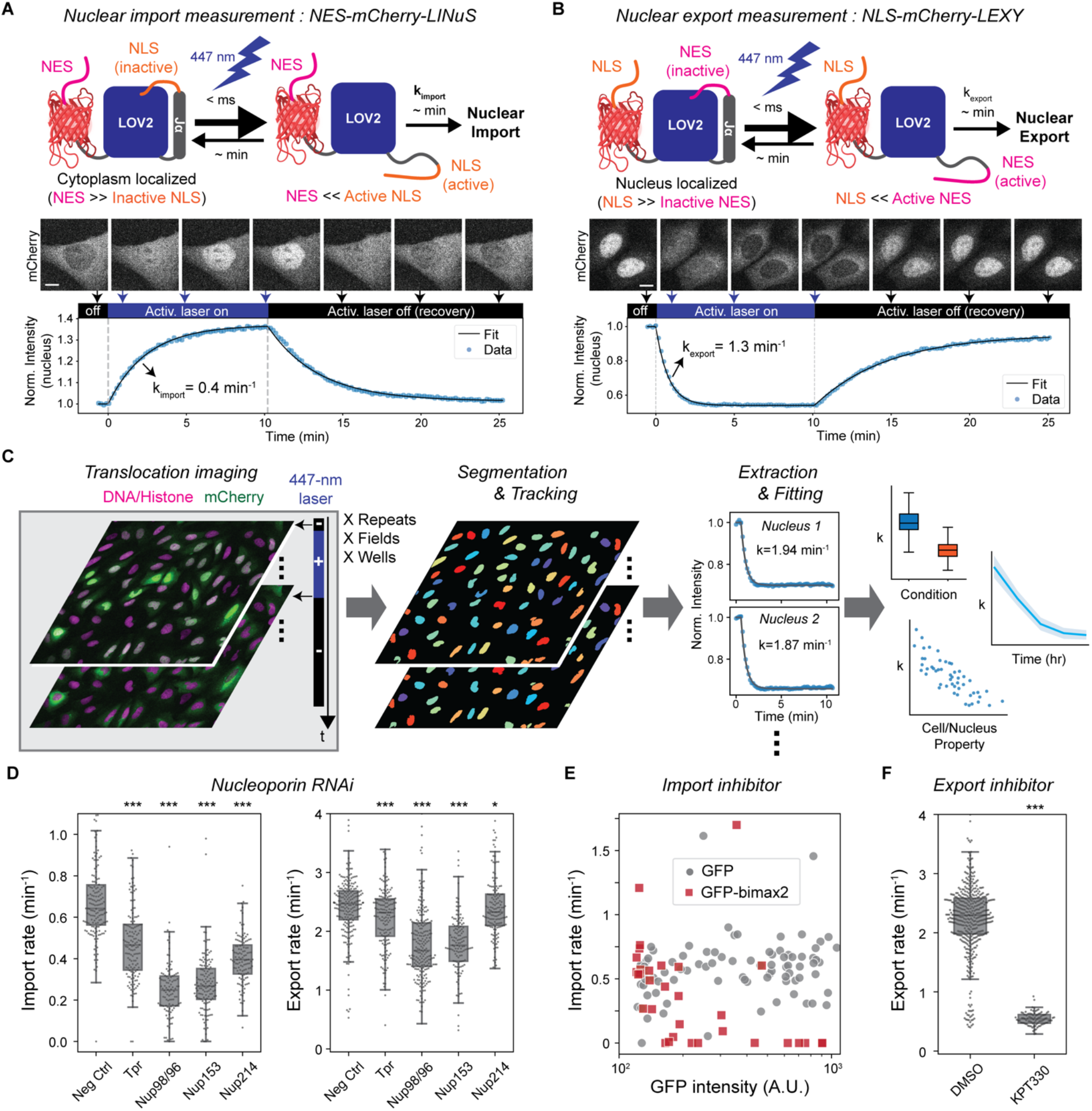
Development and validation of optogenetics-based live-cell nuclear transport assay. **(A and B)** Measurement of nuclear import and export rates using light-inducible nuclear transport systems, LINuS (import) and LEXY (export). Upon 447-nm laser illumination, NES-mCherry-LINuS and NLS-mCherry-LEXY probes translocate from the cytoplasm to the nucleus and vice versa, respectively. Nuclear transport rates are determined by fitting the light-induced change of the nuclear mCherry intensity to a mono-exponential decay model. When the activation radiation is ceased, the probes return to their pre-illumination locations, allowing repeated measurements. Scale bar, 10 μm. **(C)** Automated acquisition and analysis of nuclear transport assay. U2OS cells stably expressing the transport probes and histone-marker are imaged every 10 seconds in the absence and presence of the 447-nm activation laser illumination. The cycle of 10 min activation and 15 min recovery is repeated at multiple locations or time points, if necessary. Nuclei are segmented and tracked based on the histone or DNA images. The time course of the nuclear mCherry intensity was fitted to a mono-exponential decay model to determine the nuclear import or export rate of each nucleus. Measured transport rates were then aggregated for correlation analysis. **(D-F)** Validation of the nuclear transport assay by measuring the effects of nuclear transport perturbations: **(D)** NUP RNAi, N>100 nuclei for each RNAi condition. **(E)** GFP-bimax2 transfection for importin-αsequestration. **(F)** 30 min 1 μM KPT-330 treatment for export inhibition. N>90 nuclei. p-values were calculated by Welch’s t-test for comparison with negative control. n.s.: p>0.01; *: p<1e-2, **: p<1e-3, ***: p<1e-4. The data is also available in Table S1.

To validate the transport assays, we quantified the effects of known perturbations. We first tested the effects of siRNA-mediated depletions of the following NUPs: *NUP98*, *NUP153*, *NUP214*, and *TPR* (Fig. 1D; Fig. S1; and Table S1). Different NUP depletions differentially affected the import/export kinetics of the LINuS/LEXY probes: *NUP98* and *NUP153* depletions resulted in the largest reduction in the import rate of the LINuS probe (~60%), followed by *NUP214* (39%) and *TPR* (27%) depletions. The export rate of the LEXY probe was also greatly affected by *NUP98* and *NUP153* depletions (~30%), while slightly by *TPR* and *NUP214* depletions (~6%). These findings demonstrate the capability of the transport assays to separately measure the differential changes in the import and export rates induced by different perturbations. The differential effects of NUP depletions may reflect different roles of individual NUPs in nuclear import and export, which have been studied previously (Ball and Ullman, 2005; Bernad et al., 2006; Frosst et al., 2002; Hutten and Kehlenbach, 2006; Powers et al., 1997; Powers et al., 1995; Ullman et al., 1999; Walther et al., 2001; Wu et al., 2001).

We next tested the effect of nuclear transport inhibitors. We first inhibited nuclear import by expressing bimax2, a peptide that competitively inactivates importin-αs by tightly binding to their NLS-binding domains (Kosugi et al., 2008). We found that the import rate of the LINuS probe sharply decreased with an increasing GFP-bimax2 expression level, while GFP expression alone did not affect the rate (Fig. 1E). We next inhibited the nuclear export using KPT-330, a specific inhibitor of exportin-1 (*XPO1*), an NTR accommodating diverse NESs (Kirli et al., 2015; Lapalombella et al., 2012). After KPT-330 treatment, the export rate of the LEXY probe decreased by 75% (Fig. 1F). The import rate of the LINuS probe also decreased by 62% after KPT-330 treatment, but the relative reduction of the export rate to that of the import rate nonetheless was much greater for KPT-330 treatment than those for NUP depletions (Fig. S1; Table S1). These results demonstrate that the LINuS and LEXY probes quantitatively report on the importin-α/β1-dependent import and the exportin-1-dependent export rates, respectively, although they are not fully decoupled.

### Increasing O-GlcNAc modification accelerates nuclear import and export

With quantitative assays in hand, we investigated how nuclear transport rates respond to perturbation of O-GlcNAc modifications. We first performed siRNA transfections to deplete *OGT* or *OGA*, the sole pair of enzymes that add and remove O-GlcNAc, respectively (Chatham et al., 2020). After 3 days of the siRNA transfections, we found that the import rate was 45% lower in the OGT-depleted cells, while 11% higher in the OGA-depleted cells, compared to cells transfected with scrambled siRNA (Fig. 2A). The export rate followed the same trend as the import rate but changed to smaller extents (Fig. 2A). Consequently, the import and export rates were about 100% and 30% higher, respectively, in the OGA-depleted cells than in the OGT-depleted cells (Table S1).

We next used OGT- or OGA-specific small-molecule inhibitors, OSMI-4 (Martin et al., 2018) and Thiamet-G (Yuzwa et al., 2008), respectively, to better control the timing and degree of the OGT/OGA inhibitions. Cells were treated with each drug at three different doses and continually assayed for the nuclear transport rates over 20 hours. The import rate slightly decreased (7%) in the DMSO control over the time of on-stage incubation. It decreased significantly more in the presence of OSMI-4 in a dose-dependent manner, by up to 23% over 20 hours (Fig. 2B). The decay rate measured 0.14 ± 0.02 h^−1^, comparable to the turnover rate of O-GlcNAc at several NUPs measured previously (Wang et al., 2016). On the other hand, the import rate increased after Thiamet-G treatment in a dose-dependent manner up to 15% over 20 hours (Fig. 2B). Similarly, the nuclear export rate also reached a higher and lower level in the presence of Thiamet-G and OSMI-4, respectively, than in the DMSO control (Fig. 2B).

**Fig. 2.**
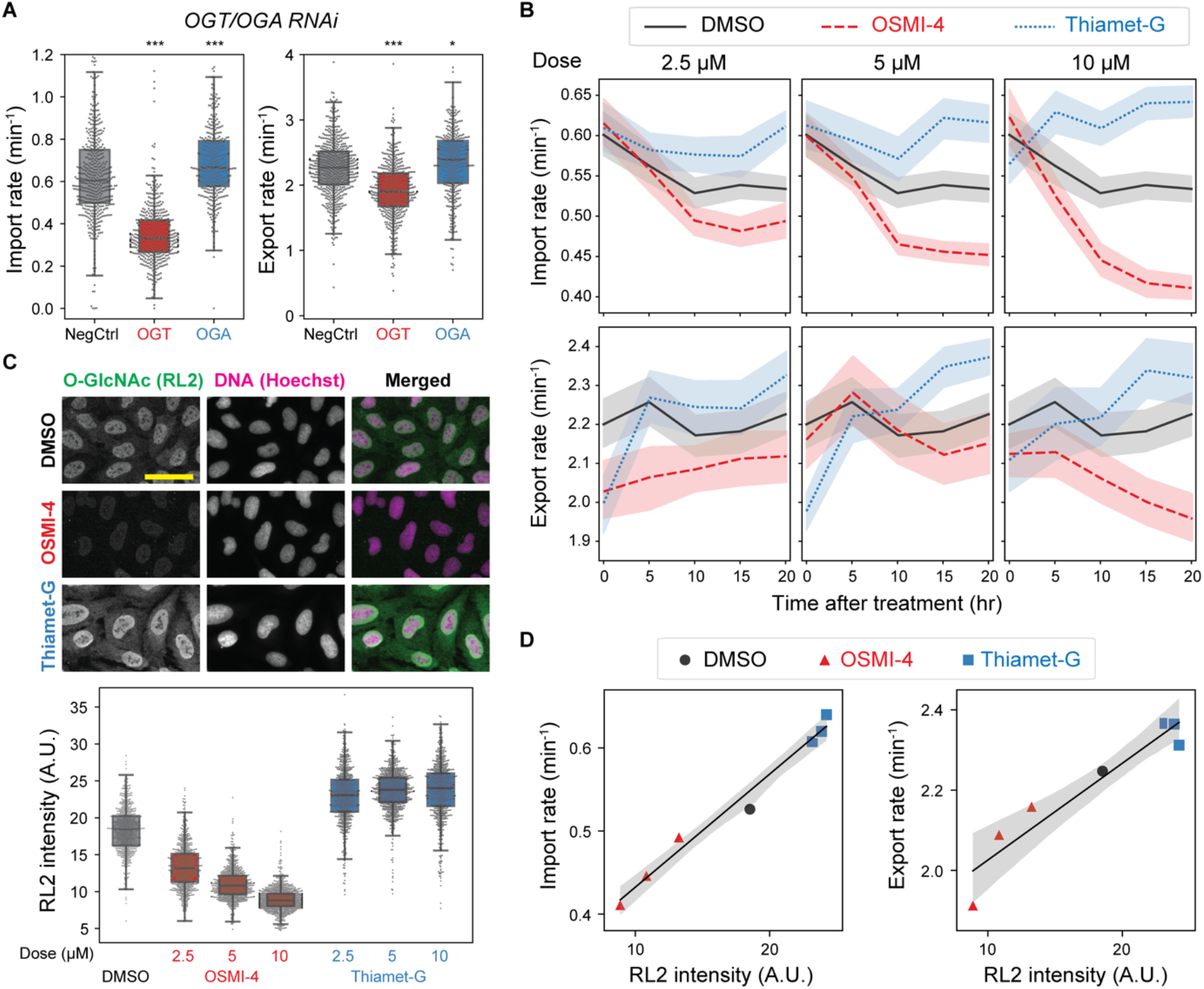
The nuclear transport rates are dependent on the O-GlcNAc level. **(A)** Nuclear import (left) and export (right) rates after 3 days of transfection with scrambled (NegCtrl), OGT or OGA siRNAs. N>360 nuclei per condition. **(B)** Import (top) and export (bottom) rates vs time after treatment with DMSO (black, solid), OSMI-4 (red, dashed) and Thiamet-G (blue, dotted) at 2.5 mM (left), 5 mM (middle), and 10 mM (right). Line and shaded area indicate mean and 95% confidence interval, respectively. N³120 nuclei per time point per condition. (**C)** After the live-cell transport assay in the presence of drugs (shown in **(B)**), cells were fixed and stained with anti-O-GlcNAc (RL2) antibody and Hoechst 33258. Representative immunofluorescence images of cells treated with DMSO (top), 10 mM OSMI-4 (middle) and 10 mM Thiamet-G (bottom). Scale bar 50 mm. Quantification of nuclear RL2 intensity (background corrected) shown in the plot below the images. N>700 nuclei per condition. (**D)** Median import (left) or export (right) rate (from **(B)**) vs median nuclear RL2 intensity (from **(C)**), measured after 20-hour treatment with DMSO (black circle), OSMI-4 (red triangle) or Thiamet-G (blue square) at three different doses. Black line and gray shade are the line fit and its 95% confidence interval. p-values were calculated by Welch’s t-test for comparison with negative control. n.s.: p>0.01; *: p<1e-2, **: p<1e-3, ***: p<1e-4. The transport assay results are also shown in Table S1.

To characterize the relationship between the O-GlcNAc level and the transport rates, we performed quantitative immunofluorescence using RL2, anti-O-GlcNAc antibody generated against glycosylated NUPs (Snow et al., 1987). Compared to the DMSO control, the nuclear RL2 signal decreased by up to 52% in the OSMI-4-treated cells and increased by up to 30% in the Thiamet-G-treated cells, exhibiting dose dependency in both directions (Fig. 2C). Plotting the transport rates (shown in Fig. 2B) against the nuclear RL2 signals (shown in Fig. 2C) revealed that both transport rates increased linearly with increasing nuclear RL2 signal, with the import rate showing a greater dependency (Fig. 2D). The relationship between the import and export rates measured after the drug-induced OGT/OGA inhibitions was colinear with that after OGT/OGA RNAi, indicating that the O-GlcNAc-dependent change in the rates is independent of the mechanism of perturbation (Fig. S1).

### Modification of NPCs drives O-GlcNAc-dependent nuclear transport modulation

We further asked whether the O-GlcNAc dependency of the transport rates was specifically due to modification of NPCs where O-GlcNAc is abundant (Davis and Blobel, 1987; Hanover et al., 1987; Holt et al., 1987; Snow et al., 1987), as opposed to less direct effects mediated by changes in gene expression (Comer and Hart, 1999) or modification of other factors in the transport machinery. Hypothetically, if the NPCs or other nucleus-confined components were responsible for the O-GlcNAc-dependent transport rates, different nuclei in a multi-nucleated cell could have different transport rates depending on their O-GlcNAc levels. To test this, we generated heterokaryons using polyethylene glycol (PEG)-mediated cell fusion (Davidson and Gerald, 1976) (Fig. 3A).

**Fig. 3.**
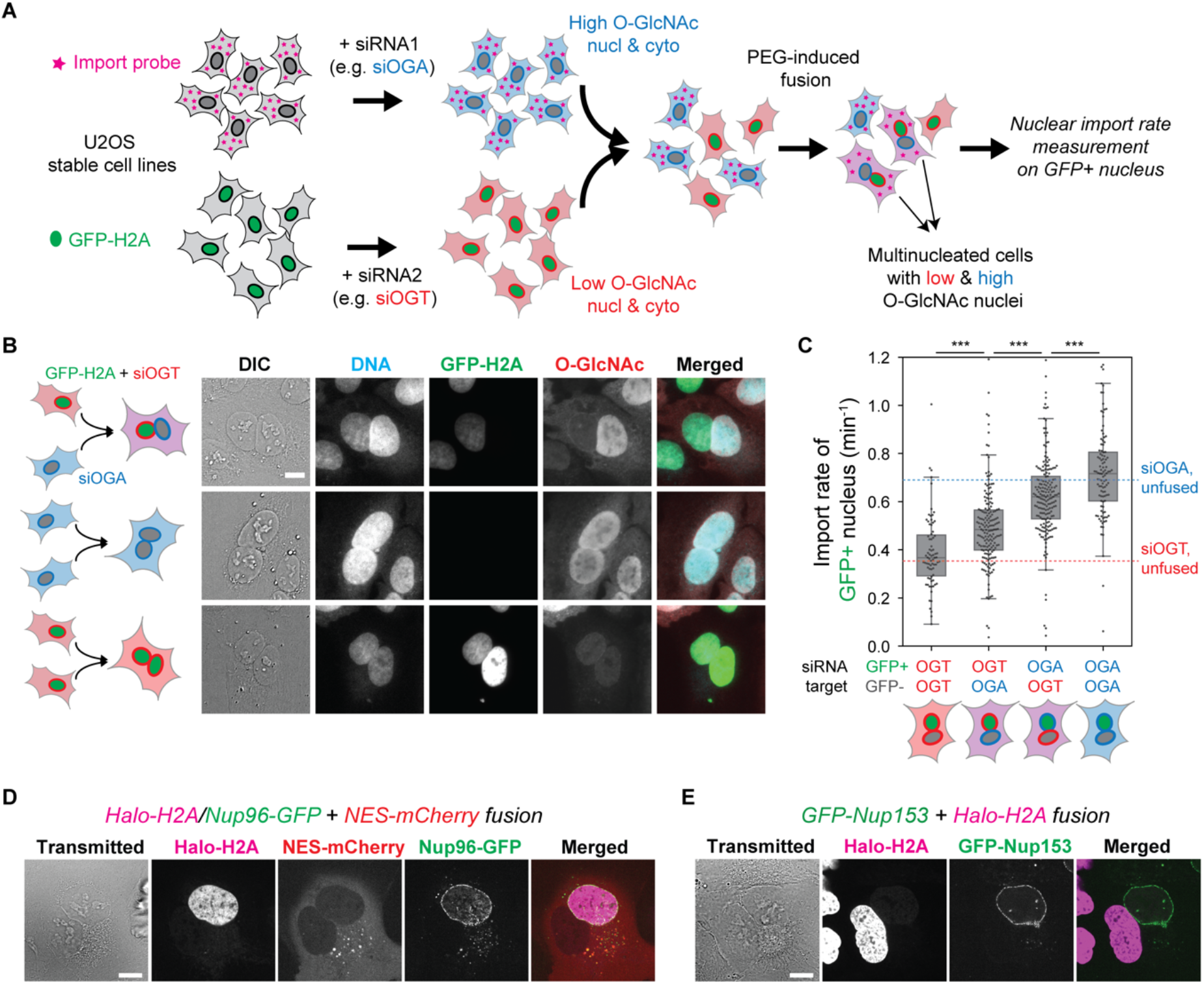
Heterokaryon analysis reveals that nucleus-confined factors are responsible for the O-GlcNAc dependency of nuclear transport kinetics. **(A)** Schematics of heterokaryon analysis. Two U2OS stable cell lines, one expressing the NES-mCherry-LINuS import probe and the other expressing GFP-H2A, were separately transfected with OGA- or OGT-targeting siRNA, co-cultured, and then fused by brief treatment with PEG. The nuclear import rate was measured on the GFP-positive nuclei in the mCherry-positive fused cells. **(B)** RL2 immunofluorescence images of the fused cells, showing that the nuclei largely retains the pre-fusion O-GlcNAc level. **(C)** The nuclear import rate of the GFP+ nucleus in four different fusions (from left to right): GFP-H2A^siOGT^+import probe^siOGT^, GFP-H2A^siOGT^+import probe^siOGA^, GFP-H2A^siOGA^+import probe^siOGT^, and GFP-H2A^siOGA^+import probe^siOGA^. Blue and red dotted lines indicate the median nuclear import rates of the unfused cells transfected with siOGA and siOGT, respectively (values from Fig. 2A). N³75 nuclei for each condition. p-values calculated by Welch’s t-test where n.s.: p>0.01; *: p<1e-2, **: p<1e-3, ***: p<1e-4. **(D)** The fusion of a U2OS cell stably expressing Halo-H2A (magenta) and Nup96-GFP (green) and another U2OS cell expressing NES-mCherry (red). **(E)** The fusion of U2OS cell stably expressing GFP-Nup153 (green) and another U2OS cell expressing Halo-H2A (magenta). Scale bars 10 mm.

Briefly, two U2OS stable cell lines, one expressing GFP-labeled histone 2A (GFP-H2A) and the other expressing the NES-mCherry-LINuS import probe, were separately transfected with OGT- or OGA-targeting siRNA. Then those two cell populations were combined and treated briefly with PEG to induce cell fusion (Fig. 3A). The multi-nucleated cells were assayed after 2-4 hours of resting in the presence of both OSMI-4 and Thiamet-G to suppress O-GlcNAc turnover. This maintained pre-fusion levels of O-GlcNAcylation in each nucleus after fusion (Fig. 3B).

We performed cell fusions with every combination of the two siRNA transfections (OGT or OGA) and the two cell lines (GFP-H2A or NES-mCherry-LINuS) and measured the nuclear import rate of the GFP+ nuclei in mCherry+ cells (Fig. 3 A and C). The GFP positivity indicates the origin of the nucleus and therefore its O-GlcNAc state, while the mCherry positivity ensures the fusion with the other origin. The siOGT/siOGT or siOGA/siOGA fusions displayed similar import rates to the OGT- and OGA-depleted unfused cells, respectively, indicating that the cell fusion itself does not alter the transport rates (Fig. 3C). Interestingly, in the siOGT/siOGA heterokaryons, the import rate was 28% higher at the high-O-GlcNAc nuclei than at the low-O-GlcNAc nuclei (Fig. 3C), indicating a significant contribution of the nucleus-confined factors to the O-GlcNAc-dependent nuclear transport regulation.

We next asked which nuclear transport machinery components are responsible for the O-GlcNAc dependency of nuclear import observed in the cell fusion assay. The NPC itself is undoubtedly the top qualified candidate not only for the abundance of O-GlcNAc modification (Davis and Blobel, 1987; Hanover et al., 1987; Holt et al., 1987; Snow et al., 1987) but also for its strict confinement to the nucleus (Daigle et al., 2001; Rabut et al., 2004). The strict nuclear confinement was confirmed by heterokaryon fusion analyses: Nup96 was not exchanged between the nuclei in a heterokaryon, neither was Nup153, despite its dynamic association with the NPC (Daigle et al., 2001; Rabut et al., 2004) (Fig. 3 D and E). We also considered the possibility that the Ran pathway plays a role in the O-GlcNAc dependency of the import rate, as Rcc1 and RanGAP1, the two major regulators of Ran (Macara, 2001), are confined to the nucleus at least partially (Mahajan et al., 1997; Matunis et al., 1996; Nemergut and Macara, 2000). However, OSMI-4 or Thiamet-G treatment did not result in any appreciable change in the distributions of Ran, Rcc1, and RanGAP1 (Fig. S2). More importantly, the nuclear/cytoplasmic ratio of RanGTP level, measured by anti-RanGTP antibody (Richards et al., 1995) immunostaining, was not affected by the O-GlcNAc perturbations, but by the overexpression of WT or mutant Ran (Ran^Q69L^ and Ran^T24N^) or Rcc1, which confirms the validity of the RanGTP immunoassay (Fig. S2). Thus, the O-GlcNAc-dependent regulation of nuclear transport rate is predominantly driven by the NPCs, rather than by the Ran-regulating system or other diffusive components of the transport machinery, such as NTRs or cargos.

### FG-repeat permeability barrier is highly modified with O-GlcNAc

To gain insight into how O-GlcNAcylation alters the NPC, we investigated the distribution of O-GlcNAc within the NPC. Previous studies showed that wheat germ agglutinin (WGA), a GlcNAc-binding lectin, is predominantly localized to the center of the NPC (Loschberger et al., 2012; Thevathasan et al., 2019). However, WGA may misrepresent the O-GlcNAc distribution due to its multiple binding sites and considerable size (36 kDa) (Nagata and Burger, 1974; Wright and Kellogg, 1996). Moreover, the O-GlcNAc distribution along the transport axis has not been revealed. We therefore employed the combination of metabolic labeling approach (Boyce et al., 2011; Woo et al., 2018; Zaro et al., 2011; Zhu et al., 2015) and copper(I)-catalyzed azide-alkyne cycloaddition (CuAAC) reaction to fluorescently label O-GlcNAc sites *in situ* and performed super-resolution microscopy (Fig. 4 A and B). To set reference points within the NPC, Nup96 was labeled with Alexa Fluor 647 via CRISPR-Cas9-mediated endogenous protein tagging and direct immunostaining using a single-domain antibody (Fig. 4C). We first imaged the double-stained cells using three-dimensional structured-illumination microscopy (3D-SIM) and confirmed a remarkably high level of O-GlcNAc at the NPCs (Fig. 4D; Video 1). We next used stochastic reconstruction microscopy (STORM) to obtain the detailed O-GlcNAc distribution within the NPC. Viewed from the top, Nup96 delineated the circular periphery of the NPC whose diameter measured 96 ± 8 nm (s.d.) (Fig. 4 E and F). O-GlcNAc was concentrated the most at the center and lower at the periphery, resembling the binding pattern of WGA (Loschberger et al., 2012; Thevathasan et al., 2019). In the side view of the NPC, Nup96 was situated along two lines delineating the top and bottom of the symmetric core of the NPC, separated by ~50 nm (Fig. 4 G and H). O-GlcNAc was localized mostly between the two lines and symmetrically distributed along the transport axis (Fig. 4 G and H). These results show that O-GlcNAc is particularly abundant within the channel of the symmetric central core of the NPC where the FG repeat permeability barrier is embedded (Lin and Hoelz, 2019).

**Fig. 4.**
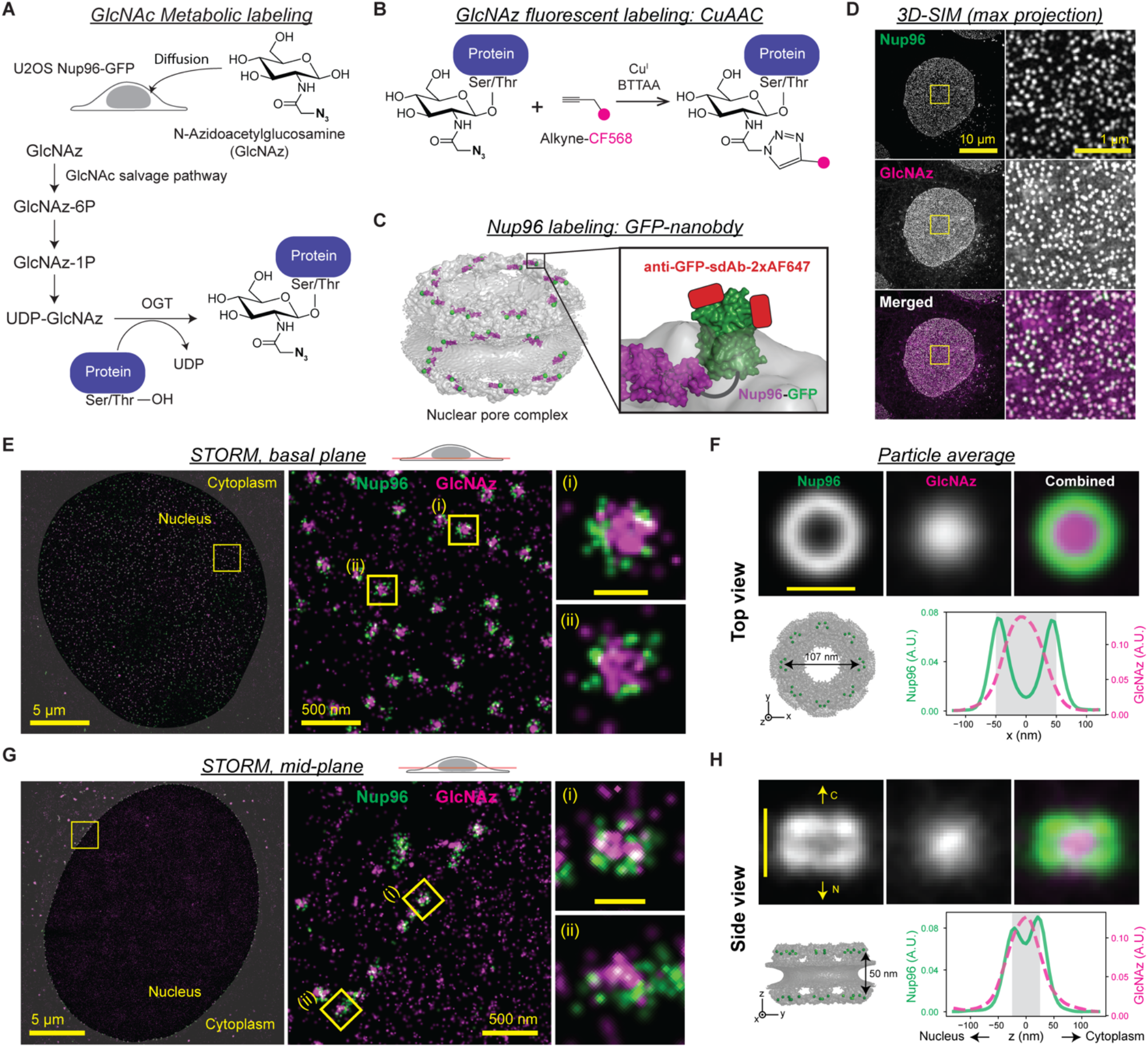
O-GlcNAc is highly abundant at the central transport channel of the symmetric core of the NPC. (**A-C)** *In situ* fluorescent labeling of Nup96 and O-GlcNAc sites in U2OS cell. **(A)** Metabolic labeling of O-GlcNAc site. GlcNAz diffused into the cells is incorporated into proteins via GlcNAc salvage pathway and OGT activity. **(B)** The incorporated GlcNAz moieties were labeled with CF568 via copper(I)-catalyzed azide-alkyne cycloaddition (CuAAC). **(C)** Nup96 (purple) was endogenously tagged with GFP (green) at the C-terminus. After fixation, each GFP molecule was labeled with up to two anti-GFP single-domain antibodies conjugated with two Alexa Fluor 647 (AF647) (red). **(D)** Maximum-intensity projection of 3D-SIM image of the double-stained cells. Left column: full field of view; right column: enlarged region of interest (yellow square). **(E and G)** Representative two-color STORM images of the basal plane (in **(E)**) and mid-plane (in **(G)**) of the nucleus showing the top and side views of the NPCs, respectively. Each localization of molecules is rendered as a normalized gaussian whose standard deviation is set to the localization precision. **(F and H)** Top: averaged top (in **(F)**) or side view (in **(H)**) STORM image of the NPCs (703 top views from 4 cells; 122 side views from 4 cells). Bottom left: structure (PDB 5A9Q) of the NPC symmetric core, where the C-terminus of Nup96 is marked with a green sphere. Bottom right: averaged intensity profiles of Nup96-GFP:AF647 (green, solid) and GlcNAz:CF568 (magenta, dashed) along the x-(in **(F)**) or z-axis (in **(H)**). Nup98-GFP:AF647 is colored in green and GlcNAz:CF568 in magenta in all merged 3D-SIM and STORM images in (**D-H)**. Scale bars in individual and averaged pore images in **(E-H)** are 100 nm.

### O-GlcNAcylation increases non-specific permeability of the FG permeability barrier

The permeability of the FG repeat barrier to the NTR-cargo complex is determined by two factors: specific binding of NTR to FG repeats and non-specific interactions between permeants and the FG barrier (Becskei and Mattaj, 2005; Bressloff and Newby, 2013; Stanley et al., 2017). O-GlcNAc modification could alter either. Previous *in vitro* studies did not report any appreciable change of NTR binding to FG domains following change in O-GlcNAcylation (Labokha et al., 2013; Tan et al., 2018). Consistent with this observation, we found that the localization of importin-β1 to NPC was unaltered after O-GlcNAc perturbations by OGT/OGA inhibitions (Fig. S3A). To determine the effect of O-GlcNAc on interactions between permeants and the barrier independent of specific bindings, we measured the transport rate of a protein probe that does not interact specifically with FG repeats. We photoconverted mEOS inside the nucleus and quantified its rate of passive permeation out of the nucleus (Fig. 5A). We found that the passive permeation rate was 22% lower in the OSMI-4-treated cells and 30% faster in the Thiamet-G-treated cells (Fig. 5A). These data followed the same trend as NTR-facilitated nuclear transport rates (Fig. 2). Since both facilitated and passive transports are accelerated by O-GlcNAcylation we conclude that it mainly increases non-specific permeation rates.

**Fig. 5.**
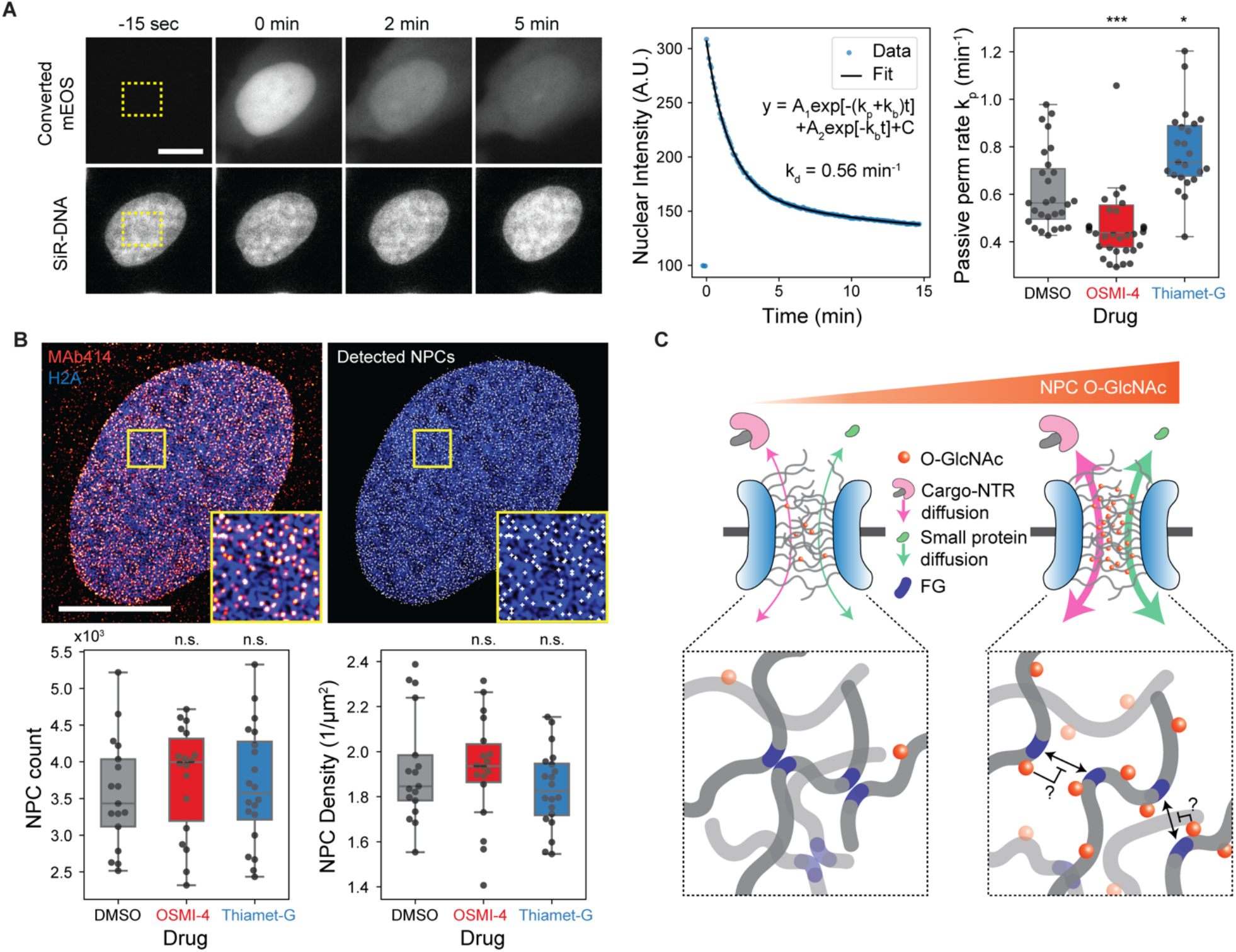
O-GlcNAcylation enhances the permeability of the NPC. **(A)** Passive diffusion rate measurement. Left representative images: mEOS (top row) in the rectangular region (yellow box) inside the nucleus was selectively converted and continually imaged along with DNA (bottom row). Middle plot: A bi-exponential decay model was fitted to the nuclear intensity of converted mEOS to measure the passive permeation rate k_p_. Right box plot: passive permeation rates for cells treated with DMSO, OSMI-4 or Thiamet-G (10 mM, 24 hour). N≥23 cells for each condition. **(B)** NPC counting. Top images: 3D-SIM image of MAb414 immunostaining (red) and Halo-H2A:JF646 (blue) and identified NPCs (white). Bottom boxplots: total number (left) and area density (right) of NPCs in cells treated with DMSO, OSMI-4 or Thiamet-G (10 mM, 24 hour). N≥17 cells for each condition. 10 mm scale bars. n.s.: p>0.01; *: p<1e-2, **: p<1e-3, ***: p<1e-4. **(C)** Proposed model for O-GlcNAc-dependent modulation of nuclear transport kinetics. Hydrophilic, bulky O-GlcNAc modifications hinder the cohesive interaction between FG repeats, thereby facilitating both passive and NTR-facilitated permeations of molecules through the NPC in both directions.

We next considered the possibility that the change in the rates does not result from the altered per-NPC permeability, but from the change in the number of NPCs, since a previous study suggested that O-GlcNAcylation protects NUPs from proteasomal degradation (Zhu et al., 2016). However, treating cells with 10 μM OSMI-4 or Thiamet-G for 24 hours did not significantly affect the number or area density of the NPCs, measured by MAb414 immunostaining and 3D-SIM (Fig. 5B; Fig. S3B). Moreover, recent proteomics data (Martin et al., 2018) of HEK293T show that the levels of NUPs, as well as other transport machinery components, do not considerably change after 24-hour treatment with 20 μM OSMI-4 (Fig. S3 C and D). Therefore, O-GlcNAcylation did not regulate NPC stability in our system. Rather, it increased non-specific permeability of the FG barrier, thereby enhancing both facilitated and passive transport in both directions (Fig. 5C). This confirms predictions made using synthetic FG-repeat-derived hydrogels (Labokha et al., 2013). The enhanced non-specific permeability might result from the hydrophilic and bulky O-GlcNAc moieties hindering the hydrophobic cohesions of FG domains (Fig. 5C).

### Concluding remarks

In summary, we developed quantitative nuclear transport assays and applied super-resolution microscopy to reveal the role of NPC O-GlcNAcylation in modulating the transport rates. The transport assays allow high-throughput quantification of the nuclear import and export kinetics in live cells, overcoming the limitations of conventional assays that rely on the measurement of the steady-state nuclear/cytoplasmic localization of cargos or on membrane-specific permeabilization of cells. The O-GlcNAc-dependent modulation of the transport rates may underlie the mechanism by which OGA inhibition mitigates the nuclear transport defects in Huntington’s disease (Grima et al., 2017) and may also provide a novel way to treat other neurodegenerative diseases and aging-related stresses compromising the nuclear transport machinery (Cho and Hetzer, 2020; D’Angelo et al., 2009; Hutten and Dormann, 2019; Kim and Taylor, 2017; Rempel et al., 2019).

## Materials and Methods

### Cell culture

U2OS cell lines (engineered from HTB-96, ATCC) were maintained in low glucose (1g/L) Dulbecco’s modified Eagle’s medium (DMEM, #10567022, Thermo Fisher) supplemented with 10% Fetal Bovine Serum (FBS, #A31605, Thermo Fisher), and 50 IU ml^−1^ penicillin and 50 μg ml^−1^ streptomycin (#15140122, Thermo Fisher), hereafter called “complete DMEM”, at 37°C in a humidified atmosphere with 5% CO_2_. Cells were validated as mycoplasma free by PCR-based mycoplasma detection kit (#30-1012K, ATCC).

### Live-cell nuclear transport assay

#### Generation of stable cell lines

U2OS cells were engineered to stably express histone H2A type 1 whose N-terminus is labeled with HaloTag (Halo-H2A) by retroviral transfection and selection with 200 μg ml^−1^ hygromycin B (#10687010, Thermo Fisher). NES-mCherry-LINuS (Niopek et al., 2014) and NLS-mCherry-LEXY (Niopek et al., 2016) coding regions in pcDNA3.1 vectors (#61347 and #72655, gift from Barbara Di Ventura & Roland Eils via Addgene) were cloned into a pBABE packaging vector for retroviral transfections by PCR and Gibson assembly. The Halo-H2A-expressing U2OS cell line was engineered to stably express NES-mCherry-LINuS (import probe) or NLS-mCherry-LEXY (export probe) by retroviral transfection and selection with 2 μg ml^−1^ blasticidin (#A1113903, Thermo Fisher).

#### Live-cell imaging

Cells were seeded at 10,000 to 15,000 per well in an 8-well chambered coverslip (#80826, ibidi) and grown in complete media for 1-3 days before live-cell imaging. Prior to imaging, the growth media was replaced with imaging media: low glucose (1g/L) DMEM without phenol red (#11054020, Thermo Fisher), supplemented with 10% FBS, 50 IU ml^−1^ penicillin and 50 μg ml^−1^ streptomycin, GlutaMAX^TM^ Supplement (#35050061, Thermo Fisher). For nucleus staining, Halo-H2A cells were incubated in 500 nM JF646-HaloTag ligand (gift from Luke Lavis) in the imaging media for 3-16 hours. Cage microscope incubator (OkoLab) was used to maintain the cells at 37 °C in 5% CO_2_ with high humidity during imaging. Live-cell imaging was performed on a Nikon Ti motorized inverted microscope with Perfect Focus System, using spinning disk confocal scanner (CSU-X1, Yokagawa) with Spectral Applied Research Aurora Borealis modification, motorized stage and shutters (Proscan II, Prior), sCMOS camera (Flash4.0 V3, Hamamatsu), laser merge module (LMM-5, Spectral Applied Research), CFI Plan Apo 20x/0.75NA objective lens (Nikon), and Chroma ZT445/514/561/640tpc polychroic mirror. mCherry-labeled import/export probes were imaged using 561 nm laser and ET605/70m emission filter (Chroma), while JF646:Halo-H2A was imaged using 642 nm laser and ET700/75m emission filter (Chroma). In the nuclear transport assays, live-cell time-lapse images were acquired through a ~25-min (155 frames) imaging cycle that consists of three acquisition phases: pre-activation, activation and recovery.

Throughout the cycle, the mCherry-labeled transport probes and DNA/histone marker were imaged every 10 seconds with 100 ms exposure time. Each imaging cycle began with a 3-frame pre-activation phase to determine the baseline nuclear localization of the probe, followed by a 10-min activation phase (61 frames) in which the probes were activated by 100 ms exposure to the activation laser (447 nm) every 10 seconds. The imaging cycle ended with 15-min recovery phase (91 frames), during which the probe returns to the pre-stimulation location in the absence of 442-nm laser stimulation, so that the imaging cycle can be repeated at the same field. To increase the throughput, the imaging cycle was executed at three different fields simultaneously. The imaging cycle was repeated at multiple time points and at different positions, which was fully automated by using “Journal” macro in MetaMorph Software. The power of 442-nm laser was optimized based on the nuclear export rate vs laser power curve, to the lowest saturation level where a small variation of the power does not influence the transport rate. No sign of photobleaching or photodamaging was noticed when the imaging cycle was repeated every hour for 24 hours at the same field.

#### Image analysis

We created custom Python codes for streamlined computational analysis of the time-lapse images. Briefly, cell nuclei were segmented based on H2A/DNA images using U-Net convolutional neural network trained on manually annotated U2OS nuclei images (Caicedo et al., 2019), and tracked using *trackpy* package (v0.4.1, 10.5281/zenodo.1226458). Then for each nucleus, the time-trajectory of the mean nuclear mCherry intensity was measured and normalized such that the average intensity during pre-activation phase is 1 and the background intensity is 0. The normalized nuclear mCherry intensity trajectory during activation phase was fit to the following mono-exponential decay model to determine the transport rate *k*:

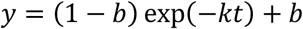

When the import probe is used, *k* corresponds to the import rate and *b* > 1 (i.e. *y* increases over time), while when the export probe is used, *k* corresponds to the export rate and 0 < *b* < 1 (i.e. *y* increases over time). Trust region reflective least-squares algorithm was used for the nonlinear regression. The fitting result was excluded from further analyses if it meets the following criteria: (1) the fitting algorithm not converged; (2) too many missing time points in the nuclear intensity trajectory (i.e. incomplete nucleus tracking); (3) reduced chi-squared statistics too large; (4) |1-b| too small; and (5) *k* abnormally large.

### siRNA transfections

siRNAs targeting Nup96/98 (SI00115332), Nup153 (SI03033226), Nup214 (SI04329521) and OGT (SI0266513) and negative control siRNA (1027280) were purchased from Qiagen, OGA siRNA (M-012805-01-0005) from Dharmacon, and Tpr siRNA from IDT (synthesized, 5’-UUUAACUGAAGUUCACCCU-3’). Cells were grown to ~70% confluency in complete DMEM for 1 day in a 6-well plate and transfected with siRNA for each well as follows: 30 pmol of siRNA and 1 μl of Lipofectamine™ RNAiMAX Transfection Reagent (#13778030, Thermo Fisher) were diluted separately in two vials of 150 μl Opti-MEM (#31985062, Thermo Fisher) and kept at room temperature for 5 minutes, while flicking once every 2 minutes. Then siRNA solution was added to the reagent solution and incubated for 30 minutes at room temperature while flicking once every 10 minutes. The growth media was replaced to Opti-MEM + 10% FBS after washing the cells with PBS (both prewarmed to 37°C), and then 300 μl siRNA/reagent cocktail was added and mixed by swirling. The transfected cells were incubated for 24 hours at 37°C/5% CO_2_, before detached and reattached onto appropriate imaging dishes for nuclear transport or cell fusion assays. The assays were performed 60-72 hours post-transfection. 80–90% depletion of OGT/OGA mRNAs was verified by RT-PCR using QuantiTect Reverse Transcription Kit (205311, Qiagen), QuantiTect SYBR Green PCR Kit (204143, Qiagen) and BioRad C1000 (base) with CFX96 (optics).

### Drug treatments

Cells were treated with 1 μM KPT-330 (#S7252, SeleckChem) for 30 minutes for exportin-1 inhibition. OGA inhibitor Thiamet-G was obtained from Cayman Chemicals (#13237), and the second-generation OGT inhibitor OSMI-4 (gift from Suzanne Walker) was obtained as described previously (Martin et al., 2018).

### Plasmid transfection

For nuclear import inhibition (Fig. 1E), psfGFP-bimax2 expression vector was constructed by inserting synthesized DNA fragment (IDT) for bimax2 sequence (RRRRRRKRKREWDDDDDPPKKRRRLD) into XhOI/EcoRI site of psfGFP-C1 vector. Cells were transfected with psfGFP-bimax2 and psfGFP-C1 vectors using TransIT-2020 transfection reagent (#MIR5404, Mirus) and imaged after 6.5 hours. For RanGTP immunofluorescence assay validation (Fig. S2C), mCherry-N1, RanWT-mRFP1 (Addgene #59750, gift from Yi Zhang), RanQ69L-mCherry (Addgene #30309, gift from Jay Brenman), RanT24N-mCherry (Addgene #37396, gift from Iain Cheeseman) and Rcc1-HA (Addgene #106938, gift from Patrizia Lavia) were transfected using TransIT-2020 24 hour before immunostaining.

### Cell fusion assay

Two different U2OS stable cell lines, one expressing GFP-H2A and the other expressing the import probe, were grown in a 6-well plate and separately transfected with siRNA targeting OGT or OGA as described above. After 24 hours, each well was washed with PBS once and treated with 300 μl of trypsin in TC incubator for ~5 min. 700 μl complete DMEM was added to each well and pipetted up and down to detach and dissociate cells. The entire 1 ml cell suspension was transferred to 1.5 ml tube, and the cell density was measured. ~10^5^ cells from each cell suspension were combined into another 1.5 ml tube, mixed by brief vortex, and centrifuged at 300g for 4 min. The supernatant was gently removed, and the cell pellet was resuspended in 1 ml complete DMEM. The resuspension was filtered through 35 μm cell strainer and transferred to a 35-mm petri dish with No. 1.5 poly-d-lysine-coated glass bottom (#P35GC-1.5-14-C, MatTek) with additional 1 ml complete DMEM. After 2 days, the co-culture was washed twice in PBS and treated with ~300 μl 50% PEG-1500 (#10783641001, Millipore Sigma) for 2 min at RT after completely aspirating the residual PBS. 3 ml PBS was gently added to the dish, swirled, and aspirated. The cells were gently washed twice more in 3 ml PBS and once in prewarmed imaging media, and then placed in 1.5 ml imaging media containing 1 μM SiR-DNA (CY-SC007, Cytoskeleton) + 10 μM Verapamil (for nucleus staining) and 10 μM OSMI-4 + 10 μM Thiamet-G (to suppress O-GlcNAc turnover). After resting in TC incubator for 2-4 hours, cells were imaged for the nuclear import measurement and analyzed as described above, or fixed and stained for immunofluorescence analysis.

### Super-resolution microscopy imaging of O-GlcNAc modification and Nup96

#### Generation of Nup96-GFP CRISPR-engineered cell line

The C-terminus of endogenous Nup96 in U2OS cells was tagged via homozygous insertion of mEGFP gene into NUP98 gene using CRISPR-Cas9 genome editing as described previously (Koch et al., 2018; Thevathasan et al., 2019). Briefly, pX335 vectors (Addgene #42355, gift from Feng Zhang) carrying SpCas9 nickase mutant (D10A) and a pair of sgRNAs, targeting GTTGGGAGCCTGTGAGCCCC and CAGTTCTCGCAGATAGGACT, were used to induce double-strand break near the C-terminal end of Nup96. Homology-directed repair donor plasmid was prepared by cloning 0.9 kb and 1 kb homology arms flanking mEGFP gene into pUC19 vector. 8-residue linker, SACYCELS, was placed between Nup96 and mEGFP. PAM sequences of the sgRNAs in the donor plasmid were muted by site-directed mutagenesis, using QuikChange site-directed mutagenesis kit (#200515, Agilent). U2OS cells were transfected with the plasmids by electroporation, using Nucleofector^TM^ 2b device (Lonza) and Ingenio^®^ electroporation kit (#50117, Mirus). After 5 days of transfection, individual GFP-positive cells were sorted into 96-well plates and expanded to monoclonal cell lines. Homozygous mEGFP insertion at the C-terminal end of Nup96 was verified by PCR.

#### Fluorescent labeling of O-GlcNAc modification and Nup96

All samples for super-resolution microscopy (presented in Fig. 4) were prepared in 35-mm petri dish with No. 1.5 poly-d-lysine-coated glass bottom (#P35GC-1.5-14-C, MatTek). For metabolic labeling of O-GlcNAc modifications, Nup96-GFP U2OS cells were grown for two days in complete DMEM containing 1 mM N-Azidoacetylglucosamine (GlcNAz, MA30911, Carbosynth), a GlcNAc analogue whose acetamido group has an azide moiety. Cells were fixed with paraformaldehyde and incubated with Alexa Fluor 647 (AF647)-conjugated anti-GFP single-domain antibody (FluoTag®-X4, #N0304-AF647, NanoTag Biotech) to fluorescently label the C-terminus of Nup96, as described previously (Thevathasan et al., 2019). Subsequently, incorporated GlcNAz was labeled with alkyne-CF568 (#92088, Biotium) via copper(I)-catalyzed azide-alkyne 1,3-dipolar cycloaddition (CuAAC) “click” reaction as follows: 100 mM CuSO_4_ and 50 mM BTTAA (#1236, Click Chemistry Tools) were premixed at 1:5 ratio. Then 500 μl of CuAAC reaction cocktail was prepared by sequentially adding 0.5 μl of 10 mM alkyne-CF568, 110 μl of CuSO_4_:BTTAA premix, and 50 μl of 1M sodium ascorbate into 340 μl of 100 mM sodium phosphate buffer pH 7 with a brief vortex after each addition. Cells were washed once in PBS with 3% BSA and then incubated in 500 μl of the CuAAC reaction cocktail for 30 min at room temperature while being protected from light. Sample was then washed twice in PBS with 3% BSA and twice in PBS for 10 min each. GlcNAc/Nup96 double-stained samples were stored in a sealed petri-dish covered with aluminum foil at 4°C for less than a month before being imaged.

#### 3D structured-illumination microscopy (3D-SIM)

Buffer was exchanged to a mounting media (90% glycerol, 0.5% propyl-gallate, 20mM Tris-HCl pH 8.0) and sealed. 3D-SIM images were acquired on GE DeltaVision OMX Blaze with Olympus 60X/1.42NA Plan Apo oil immersion objective and front-illuminated scientific-CMOS camera (PCO). For GlcNAz:CF568 imaging, 568-nm excitation laser, 571/19 (center wavelength/bandwidth) excitation filter, and 609/37 emission filter were used. For Nup96:AF647 imaging, 642-nm excitation laser, 645.5/15 excitation filter, and 683/40 emission filter were used. 40-60 optical sections were imaged every 125 nm for each channel. 3D-SIM reconstruction was performed using a CUDA-accelerated version of the conventional algorithm (Gustafsson et al., 2008), using measured, wavelength-specific OTFs matched to the data (code available at https://github.com/scopetools/cudasirecon). The final reconstructed 3D-SIM images have 40.96 μm X 40.96 μm (1024 px X 1024 px) field of view and 135-160 nm lateral and 350-380 nm axial resolutions. Channel registration was achieved via maximization of cross-correlation.

#### Stochastic reconstruction microscopy (STORM) setup

STORM data were acquired on Nikon Ti motorized inverted microscope with Perfect Focus System, equipped with Nikon TIRF illuminator and Prior Proscan II motorized stage, filter wheels and shutters. For illumination, we used the output of Agilent MLC400B monolithic laser combiner with 405 nm, 561 nm and 647 nm laser lines, which was reflected by quad-band dichroic beamsplitter, ZT405/488/561/647rpc (Chroma). Emitted fluorescence was collected by CFI Apo TIRF 100x/1.49NA oil immersion objective (Nikon) and filtered by the aforementioned dichroic beamsplitter, additional quad-band emission filter (ZET405/488/561/647m, Chroma), and an emission filter ET600/50m (Chroma) or ET700/75m (Chroma) for CF568 and AF647 emissions, respectively. The filtered emission was projected onto EMCCD camera (iXon Ultra DU-897U-CS0-#BV). Additional 1.5x intermediate magnification was applied, which results in a pixel size of 104.3 nm. NIS-Elements software was used to control the hardware.

#### STORM acquisition

We used GLOX/MEA/COT STORM buffer, which is 86.5 mM Tris-HCl pH 8, 8.65 mM NaCl, 13.65% (w/v) glucose, 560 μg/ml glucose oxidase (#G2133, Millipore Sigma), 40 μg/ml catalase (#C100, Millipore Sigma), 35 mM cysteamine (MEA, #30070, Millipore Sigma) and 2 mM cyclooctatetraene (COT, #138924, Millipore Sigma). The buffer was prepared fresh by mixing stock solutions right before imaging and replaced every 90 minutes. Sample in 35-mm glass-bottom dish was briefly washed once in 0.5 ml STORM buffer, and then placed in 1.5 ml STORM buffer and clipped down on the microscope stage. Nup96-GFP-AF647 was live-imaged briefly with weak (0.5%) 647-nm illumination to adjust xy position and focus as well as TIRF angle to achieve highly inclined and laminated optical sheet (HILO) illumination (Tokunaga et al., 2008). The focus was set to either basal plane or mid-plane of the nucleus so as to acquire top and side views of NPCs, respectively. STORM data for Nup96:AF647 and GlcNAz:CF568 were acquired sequentially using maximum-power 647-nm and 568-nm laser, respectively, while the power of 405-nm stimulation was being modulated constantly so as to have 20-40 molecules per frame. ~6000 and 40000 frames were imaged for Nup96:AF647 and GlcNAz:CF568, respectively, with 30 ms exposure time. Camera was set to readout rate 10 MHz, pre-amplifier setting 1, EM gain 300x throughout imaging.

#### STORM post-processing

STORM image reconstruction was performed using ImageJ plugin ThunderSTORM (Ovesny et al., 2014) as follows: Difference of Gaussians filter with standard deviations 1 and 1.6 pixels and local maximum detector were used to identify single molecule spots. Each spot was fitted with a symmetric Gaussian PSF model using maximum likelihood estimation (MLE). The xy positions were corrected for residual drift by redundant cross-correlation-based drift correction method. Localizations were filtered by the localization precision (<10 nm) and the size of Gaussian PSF (sigma<150 nm). Localizations that appears in consecutive frames (with a maximum gap of one missing frame) within 30 nm from one another were merged into one localization. Each single-molecule coordinate was rendered as a normalized 2D Gaussian whose standard deviation is equal to the localization precision, where the pixel size of the reconstructed images was set to 1/20 of the raw image pixel size, i.e. 104.3/20 = 5.215 nm. Reconstructed STORM images of Nup96:AF647 and GlcNAz:CF568 were registered by a custom algorithm written in Python. Briefly, 2D histogram of single-molecule localizations was constructed for each channel where the bin size was set to 20 nm x 20 nm. Then cross-correlation between two 2D histograms was calculated, and then the peak of the cross-correlation was localized to a sub-bin accuracy by fitting the peak to a symmetric 2D Gaussian using Levenberg-Marquardt least squares optimization. Finally, Nup96:AF647 STORM image was translated by the offset of the cross-correlation peak from the center.

#### Particle averaging

The top view average of the NPC was computed using a custom Python code as follows: NPC particles were detected based on the basal nuclear plane STORM image of AF647, by applying Gaussian blurring and detecting the local maxima. Anomalous particles were filtered out based on the peak CF568 and AF647 intensities. xy positions of AF647 within the 240 nm X 240 nm window were fit to a circle using robust least squares algorithm, which minimizes 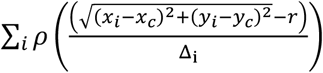, where (*x*_*c*_, *y*_*c*_) and *r* are the center position and radius of the circle, Δ_i_ the localization precision of i-th AF647 position (*x*_*i*_, *y*_*i*_), and 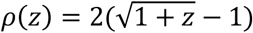 the soft l1 loss function. The low-quality NPC particles were further filtered out based on the following exclusion criteria: (1) *r* < 40 nm or r > 60 nm; (2) standard errors of *r*, *x*_*c*_, *y*_*c*_ < 5 nm; (3) number of AF647 localizations in the window less than 15. The 240 nm X 240 nm STORM images of the remaining NPC particles were translationally aligned based on the center positions (*x*_*c*_, *y*_*c*_) of the fitted circles and averaged. The side view average was obtained using the mid-plane STORM images, by manually determining the center and orientation of the NPCs using Fiji software and translationally and rotationally aligning the NPCs using a custom Python code. No cell-to-cell variation in the averaged images was observed.

### Immunofluorescence

#### Sample preparation

All immunofluorescence samples except those for super-resolution microscopy shown in Fig 4 were prepared as follows: Cells were fixed using 4% PFA/PBS for 10 min, permeabilized using 0.1% Triton-X/PBS for 10 min, and blocked using 1% BSA/PBS for 1 hour. Primary antibodies were diluted using the following ratios in 1% BSA/PBS: anti-O-GlcNAc (RL2, 1:200, #ab2739, Abcam), anti-Rcc1 (1:50, #3589S, Cell Signaling), anti-Ran (1:800, #ab53775, Abcam), anti-RanGAP1 (1F3A4, 1:100, #67146-1-Ig, Proteintech), anti-FG-NUPs (MAb414, 1:1000, #902901, Biolegend), anti-RanGTP (AR12, 1:200, gift from Ian Macara), anti-Importin-β1 (3E9, 1:1000, #MA3-070, Thermo Fisher), anti-mCherry (8C5.5, 1:500, #677701, Biolegend), and anti-HA (16B12, 1:1000, #901516, Biolegend). Cells were incubated with the diluted primary antibodies for 1 hour at RT or overnight at 4°C in a humidified chamber. The cells were incubated with goat anti-rabbit or anti-mouse secondary antibody conjugated with Alexa Fluor 488 or Alexa Fluor 568 (#A11008/#A11001/#A11031, Thermo Fisher) diluted (1:1000 ratio) in 1% BSA/PBS for 1 hour at RT. Cells were washed in PBS for 5 minutes three times after each incubation step. GFP was detected by using Alexa Fluor 647-conjugated anti-GFP single-domain antibody (FluoTag®-X4, #N0304-AF647, NanoTag Biotech) at 1:500 dilution for 1 hour at RT. SiR-DNA (#CY-SC007, Cytoskeleton), JF646-HaloTag ligand (gift from Luke Lavis), or Hoechst 33258 (#09460, Polyscience) were used for nuclei counterstaining.

#### Image acquisition

Immunofluorescence images shown in Fig. 2, Fig. 3 and Fig. S3 were acquired using Nikon Ti motorized inverted microscope equipped with Perfect Focus System, Yokagawa CSU-X1 spinning, Spectral Applied Research LMM-5 laser merge module, and Hamamatsu ORCA-R2 cooled CCD camera. CFI Plan Apo 20x/0.75NA objective lens (Nikon) was used for Fig. 2 and Fig. 3, while CFI Plan Apo Lambda 100x/1.45NA Oil objective lens (Nikon) was used for NPC localization analysis of Importin-β1 (Fig. S3A). 488 nm laser and ET525/50m filter (Chroma) were used for AF488 imaging, 561 nm laser and ET620/60m filter (Chroma) for AF568, and 642 nm laser and ET700/75m filter (Chroma) for AF647 and JF646. Di01-T405/488/568/647 beam splitter (Semrock) was commonly used for all channels. MetaMorph Software was used to control the microscope. Immunofluorescence images shown in Fig. S2 were acquired using Nikon Eclipse Ti2 motorized inverted microscope equipped with Perfect Focus System, Lumencor sola light engine, CFI Plan Apochromat Lambda 20x/NA0.75 objective lens (Nikon), and Andor Zyla 4.2 sCMOS. FITC filter set (ET470/40x ex, ET525/50m em, T495lpxr), TRITC filter set (ET545/25x ex, ET605/70m em, T565lpxr mirror), and Cy5 filter set (ET640/30x ex, ET690/50m em, T660lpxr mirror) were used for AF488, 568 and 647 imaging, respectively. NIS-Elements software was used to control the hardware.

#### Image analysis

U-Net convolutional neural network (described above) or *ilastik* (Berg et al., 2019) was used for nucleus and cytoplasm segmentations. Importin-β1 localization analysis (Fig. 3A) was performed as follows: Each of Nup96-GFP:AF647 and importin-β1:AF568 images was subtracted by the cytoplasmic background level that was determined by finding the mode intensity value of the gaussian-blurred image (sigma = 2 pixels).

Nuclear mask was generated based on the Nup96 image by gaussian blurring, thresholding and removal of small holes. The background-subtracted pixel intensities of importin-β1 vs Nup96 within the nuclear mask were fitted to a linear model by robust least squares fitting algorithm using soft l1 loss. The slope of the linear model was used as a measure of the NPC localization of importin-β1.

### Photoactivation assay for passive permeation rate measurement

U2OS cells were transfected with mEOS4b-C1 vector (Addgene #54812, gift from Michael Davidson) using Nucleofector^TM^ 2b device (Lonza) and Ingenio^®^ electroporation kit (#50117, Mirus), and seeded onto a 35-mm petri dish with No. 1.5 poly-d-lysine-coated glass bottom (#P35GC-1.5-14-C, MatTek). After resting for 4-6 hours in the complete DMEM in the TC incubator, cells were incubated in the complete DMEM containing 10 μM OSMI-4, 10 μM Thiamet-G or DMSO for 22 hours. 1-3 hours before imaging, the drug-containing complete DMEM was replaced by the imaging media containing the same drug and 500 nM SiR-DNA (CY-SC007, Cytoskeleton) + 5 μM Verapamil. Photoactivation assay was performed on GE DeltaVision OMX Blaze, the same instrument used for 3D-SIM and described above, while the sample was maintained at 37°C by stage-top heater and objective heater. 568-nm laser, 571/19 (center wavelength/bandwidth) excitation filter and 609/37 emission filter were used for imaging converted mEOS while 642-nm laser, 645.5/15 excitation filter and 683/40 emission filter were used for imaging SiR-DNA. Images were taken every 5 seconds for 15 minutes with 50 ms exposure time. After the 3rd time point, 0.5-sec pulse of 100% power 405-nm laser was shone onto a rectangular region inside the nucleus by high speed galvo in order to selectively convert mEOS in the nucleus. The time-course of the nuclear intensity of the converted mEOS was fitted to a bi-exponential model, *A*_1_ exp[−(*k*_*p*_ + *k*_*b*_)] + *A*_2_ exp[−*k*_*b*_] + *C*, where *k*_*p*_ and *k*_*b*_ are the passive permeation rate and the bleaching rate, respectively. We did not observe any significant difference in the bleaching rate between different O-GlcNAc perturbation conditions (data not shown).

### Statistical analysis

All reported point estimations are median or calculated based the median values (e.g. percentage change in nuclear transport rates) unless stated otherwise. All reported p-values were calculated using Welch’s t-test, after removing outliers that are higher than Q3 + 1.5IQR or lower than Q1 – 1.5IQR, where Q1 and Q3 are the 1st and 3rd quartiles and IQR (inter-quartile range) is Q3-Q1. Batch effect was examined for every data, which was seen for the data shown in Fig. S2B, Fig. S3 A and B. For those data, each batch was normalized such that the median of the negative control (e.g. DMSO-treated cells) is equal to 1.

## Supplemental materials

Fig. S1 and Table S1 summarize the results of nuclear import and export rate measurements performed in this study. Fig. S2 shows the nuclear and cytoplasmic localizations of Ran, Rcc1, RanGAP1, and RanGTP measured after O-GlcNAc perturbations. Fig. S3. shows importin-β1 localization to NPCs and protein levels of nuclear transport machinery after O-GlcNAc perturbations. Videos 1 shows 3D z-stack of Nup96-GFP:AF647 (green, left) and GlcNAz:CF568 (magenta, right). Z-spacing 130 nm.

## Acknowledgements

This study was supported by the NIH/NIGMS research grant R35GM131753 and postdoctoral fellowship F32GM131585. We thank Prof. Suzanne Walker, Dr. Zebulon Levine and George Fei (Harvard Medical School) for providing OSMI-4 and RL2 antibody and advice on O-GlcNAc experiments; Prof. Ian Macara (Vanderbilt University) for providing anti-RanGTP antibody; Dr. Luke Lavis (Janelia Research Campus) for providing Janelia Fluor dyes; and Dr. Yuyu Song (Mass General Hospital) for providing anti-Ran antibody. We also thank the Nikon Imaging Center and Cell Biology Microscopy Facility at Harvard Medical School, especially Dr. Talley Lambert, for help with light microscopy and comments on the data and method section.

The authors declare no competing interests.

## Author contributions

T.Y.Y. and T.J.M. conceived the study and wrote the manuscript. T.Y.Y. designed and performed the experiments and analyzed the data with input from T.J.M.

