## Supplemental Figures for "O-GlcNAc modification of nuclear pore complexes accelerates bi-directional transport"

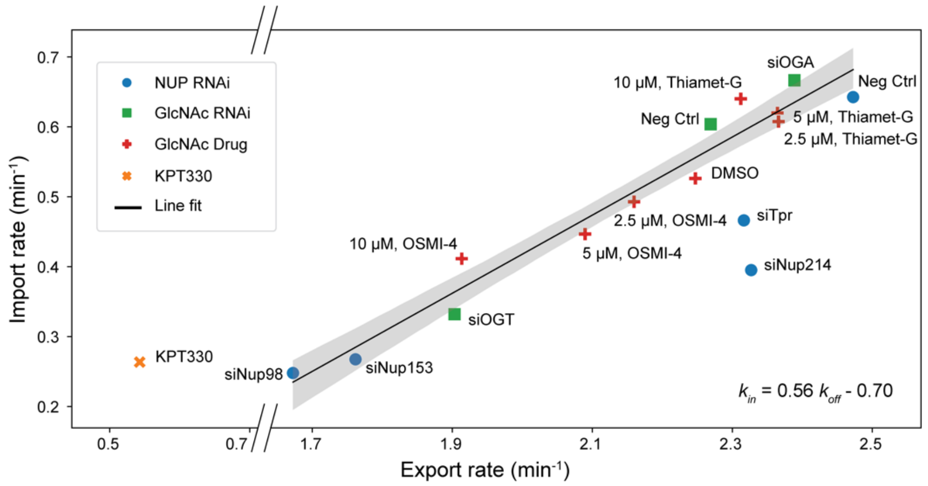


**Fig. S1.** Nuclear import and export rates measured after NUP depletions (blue dots), OGT/OGA depletions (green squares), OGT/OGA inhibitor treatments (red crosses), and KPT-330 treatment (orange x-cross). Black line represents the best line fit to the all data points excluding three outliers, siTpr, siNup214 and KPT-330, and shaded area represents the 95% confidence interval.

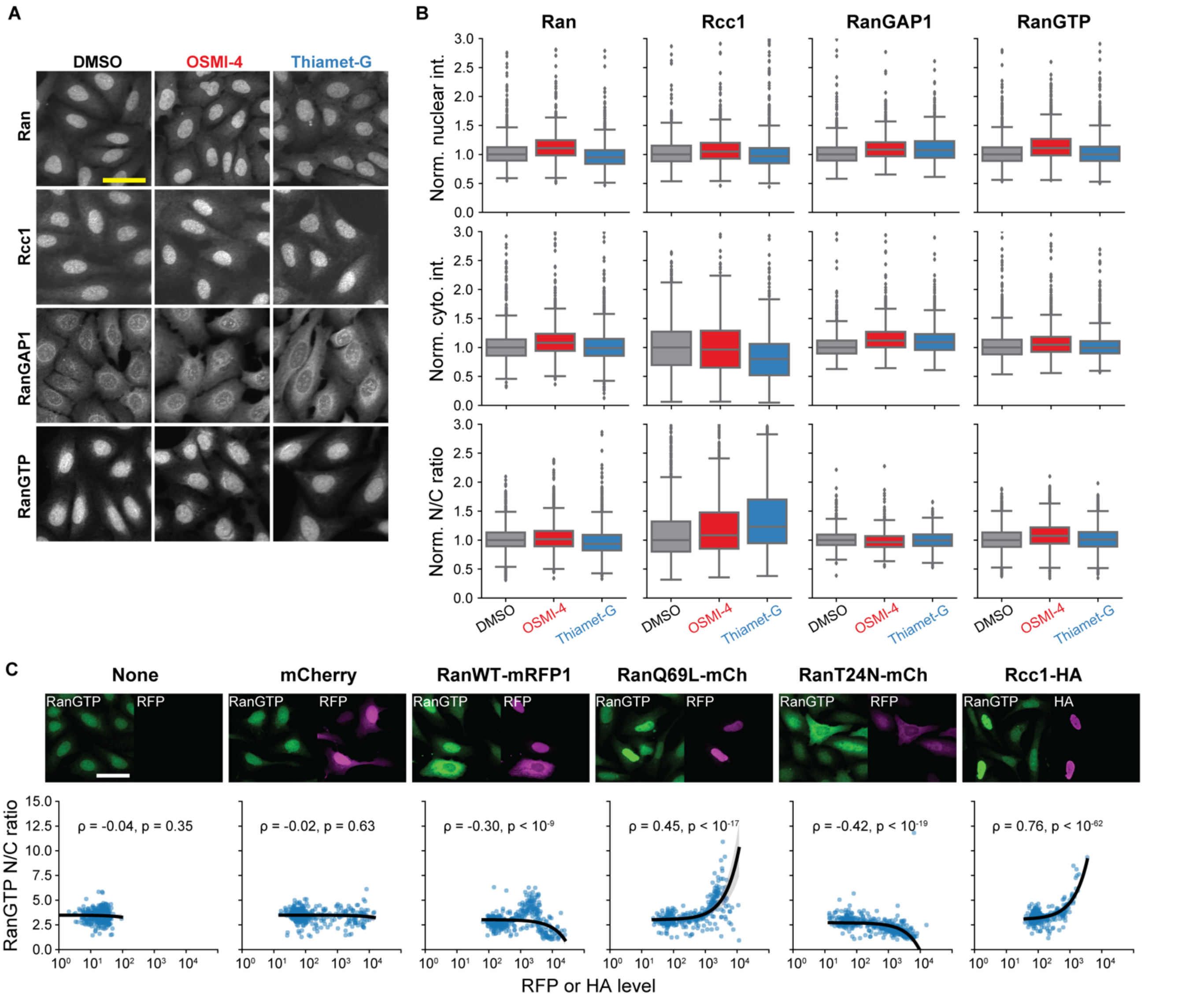


**Fig. S2. RanGTP gradient is unaltered by O-GlcNAc perturbations. (A)** Ran, Rcc1, RanGAP1 and RanGTP immunofluorescence images of cells treated with DMSO, 10 µM OSMI-4 or 10 µM Thiamet-G for 24 hours. **(B)** Mean nuclear and cytoplasmic intensity and the nucleus-to-cytoplasm intensity ratio of Ran, Rcc1, RanGAP1 and RanGTP. Data from the same batch was normalized to the median value of the DMSO condition. N>1400 cells for each condition. **(C)** (top) RanGTP immunofluorescence images of cells transfected with mCherry, RanWT-mRFP1, Ran69L-mCherry, RanT24N-mCherry, or Rcc1-HA. (bottom) Scatter plots of RanGTP nucleus-to-cytoplasm intensity ratio vs RFP or HA level. N>300 cells for each condition. Black line is the line fit to the data. Shaded gray area represents the 95% confidence interval. Scale bars 50 µm. ρ: Pearson correlation coefficient. p: p-value.

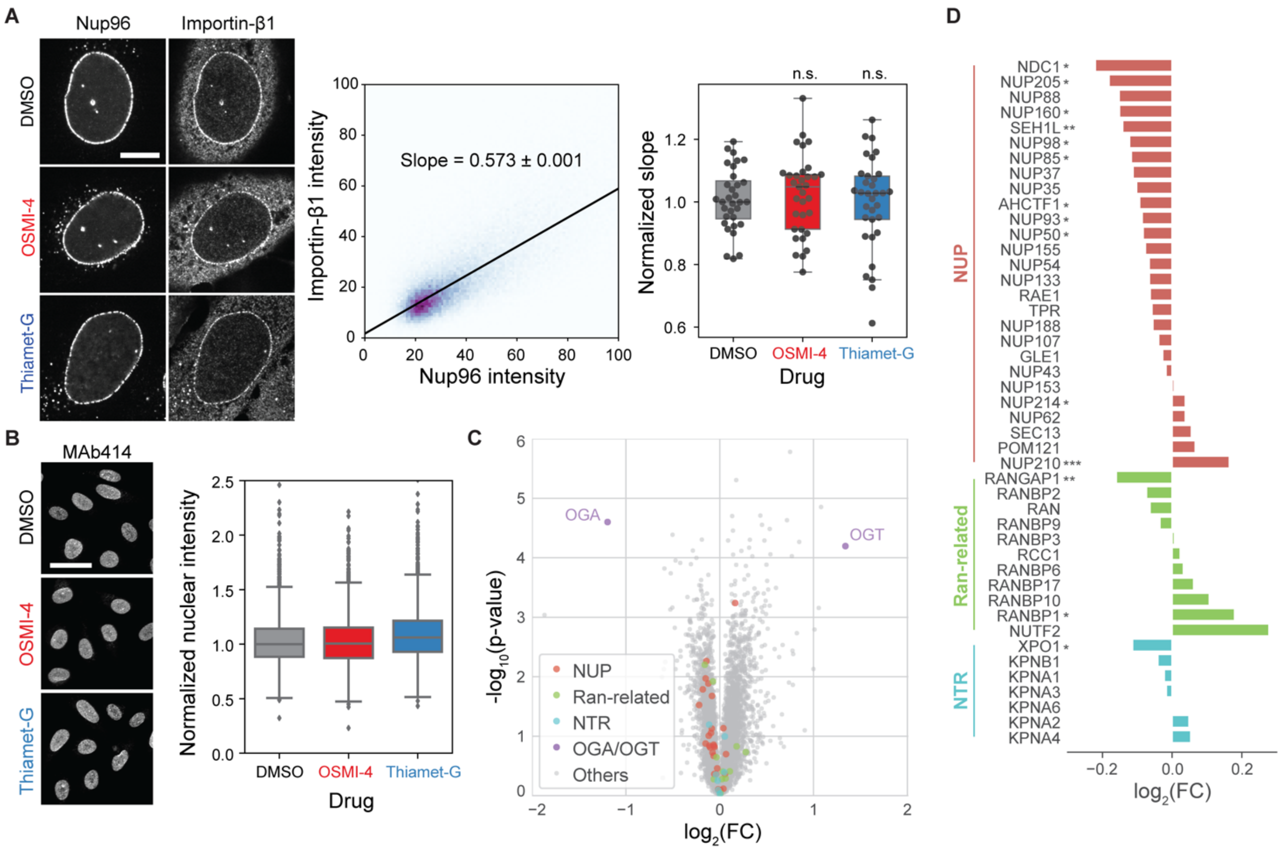


**Fig. S3. O-GlcNAc perturbations do not affect importin-β1 localization to NPCs and protein levels of nuclear transport machinery. (A)** (left) Importin-β1 and Nup96-GFP immunofluorescence images of cells treated with DMSO, 10 μM OSMI-4 or 10 μM Thiamet-G for 24 hours. (middle) For each cell, a linear model (black line) was fitted to the pixel intensities of Importin-β1 and Nup96 after background subtraction and thresholding. The slope of the linear model was used as a measure of the localization of Importin-β1 at the NPCs. (right) The slopes for each drug condition. N>31 cells for each condition. n.s.: p>0.5. **(B)** (left) MAb414 immunofluorescence images of cells treated with DMSO, 10 μM OSMI-4 or 10 μM Thiamet-G for 24 hours and (right) quantified mean nuclear intensity. N>2000 cells for each condition. For both **(A)** and **(B)**, data from the same batch was normalized to the median value of the DMSO condition. **(C and D)** Proteomics data from Martin *et al*. replotted to show the difference in the protein levels of nuclear transport machinery components between HEK293T cells treated with DMSO and those treated with 20 μM OSMI-4 for 24 hours. *: p<0.1, **: p<0.01, ***: p<0.001.
